# Replication stress in activated human NK cells induces sensitivity to apoptosis

**DOI:** 10.1101/2023.09.07.556586

**Authors:** Nicole C. Guilz, Yong-Oon Ahn, Hijab Fatima, Luis Alberto Pedroza, Seungmae Seo, Rajesh Kumar Soni, Ning Wang, Dieter Egli, Emily M. Mace

## Abstract

Natural killer cells are innate immune effectors that kill virally infected or malignant cells. Natural killer cell deficiency (NKD) occurs when NK cell development or function are impaired, and individuals with NKD are susceptible to severe and recurrent viral infections. Several gene deficiencies result in NKD, including variants in MCM4, GINS1, MCM10 and GINS4, which are components of the CDC45-MCM-GINS (CMG) helicase. The CMG helicase unwinds DNA during replication and is expressed in any actively proliferating cell. NK cells are more strongly impacted by mutational deficiencies in helicase proteins than other lymphocytes, though the mechanisms underlying this susceptibility are not completely understood. NK cells from individuals with NKD as a result of helicase deficiency have increased DNA damage, cell cycle arrest, and replication stress. Here, we induced replication stress in activated mature NK cells or T cells by chemical methods, using aphidicolin, and through shRNA knockdown of MCM10 in an NK cell line. We found that the CD56^bright^ subset of NK cells accumulates more DNA damage and replication stress during activation than CD56^dim^ NK cells or activated T cells. Aphidicolin treatment increases apoptosis of CD56^bright^ NK cells through increased pan-caspase expression and decreases perforin expression in surviving cells. This effect is modeled by shRNA mediated knockdown of MCM10, thus linking decreased helicase protein expression to replication stress and impaired NK cell function. These findings show that sensitivity to replication stress affects human NK cell survival and function and can contribute to NK cell deficiency.

## Introduction

Natural killer cells are innate lymphocytes that lyse virally infected or malignant cells without prior sensitization through germline activating and inhibitory receptors. Human peripheral blood NK cells are present in two subsets; mature CD56^dim^ cells which make up the majority of circulating NK cells and have greater baseline cytotoxic function, and less mature CD56^bright^ subset, which are potent cytokine producers (1, 2). Despite their lack of germline-encoded antigen specific receptors, NK cells can also have features of immune memory, including rapid expansion upon secondary infection, and epigenetic and transcriptional changes that lead to the generation of long-lived memory cells (3–6). Human cytomegalovirus (HCMV) infection can induce the production of memory NK cells through NKG2C-HLA-E interactions (3, 7–9). Other viruses, such as EBV and HIV, induce a subset of long-lived memory NK cells with increased effector functions upon recall (3). Adaptive NK cells can be modeled in vitro using cytokines, termed cytokine induced memory-like (CIML) NK cells, which occurs following stimulation with IL-15, IL-18, and IL-12 and results in NK cell activation of mouse and primary human NK cells (10–12). IL-15 is critical for NK cell maturation and survival, while IL-18 and IL-12 are produced by antigen presenting cells (APCs) in response to viral infection (13, 14). Thus, CIML NK cells proliferate and upregulate cytotoxic responses upon stimulation and can serve as a useful model of primary NK cell activation during viral infection. In vivo, these activated NK cells exhibit heightened responses against infections and cancerous target cells (12, 15, 16).

The absence of NK cells, dysregulation of NK cell maturation, or impairment of NK cell function leads to natural killer cell deficiency (NKD), an inborn error of immunity (IEI) that is associated with recurrent viral infections or cancer (17, 18). Though rare, partial loss-of-function variants in MCM4, GINS1, MCM10, and GINS4 lead to immunodeficiency marked by severe and recurrent viral infections and impaired NK cell maturation and function (19–23). Partial loss-of-function in these contexts is generally defined by decreased protein expression or function while residual activity is maintained. Indeed, total knockout of helicase genes leads to embryonic lethality in multiple mouse models (24–26). These genes encode components of the CDC45-MCM-GINS (CMG) helicase complex, which is required for DNA replication in S phase by unwinding DNA and recruiting polymerases (27, 28). The CMG helicase is made up of a core of 11 subunits including CDC45, MCM2-7 and GINS1-4 with additional proteins including MCM10. The helicase is active in any proliferative cell and plays an important role in maintaining genomic stability (27, 28). Individuals with NKD due to partial loss-of-function of helicase proteins have increased DNA damage and apoptosis, cell cycle arrest, and replication stress in NK cells and other cell subsets, however the mechanism behind the accumulating evidence that NK cells are more sensitive than other lymphocytes or proliferating cells is poorly understood (19–23, 29).

Replication stress occurs when a replication fork is stalled or traveling too slow for proper function (30). Stress at the replication fork arises from endogenous sources includ-ing DNA lesions such as inter-strand crosslinks (ICLs), dormant replication origins, and erroneous incorporation of nucleotides during replication in S phase (30). Dysregulation of CMG helicase proteins leads to replication stress. Over-expression of Cdc45 in Xenopus extracts results in replication stress through increased origin firing and density, and increased asymmetrical replication forks (31). The same study overexpressed the GINS complex and found similar results with decreased inter-origin distance and similarly increased fork asymmetry (31). Transient knockdown of MCM3 and MCM2 with siRNA similarly resulted in replication stress as evidenced by DNA damage and activation of the DNA damage response (DDR) pathway (32). RNAi against MCM5 developed increased sensitivity to replication stress from chemical inhibition of replication with an accumulation of DNA damage and cell cycle arrest phenotypes (33). The CMG helicase complex and associated proteins also serve as molecular protectors against endogenous causes of replication stress. MCM10, which activates origins and maintains stability during elongation, can balance replication stress in cancer cells by activating dormant origins (34–36). Thus, loss of helicase function results in replication stress. However, NK cells are particularly susceptible to helicase protein deficiency while other lymphocytes such as CD3+ T cells appear to exhibit tolerance. Helicase deficiency may have an impact on the development of NK cells, such as blocking the maturation into the CD56^dim^ subset (19–21, 23). However, while helicase deficiencies have been shown to affect NK cell development, the ubiquitous expression of helicase proteins led us to predict that partial loss-of-function variants can also affect mature NK cell function. We thus sought to investigate whether the unusual susceptibility of NK cells to replication stress induced by helicase deficiencies affects mature NK cell function in addition to NK cell maturation.

Here, we demonstrate that mature NK cell subsets are more sensitive to replication stress than T cells during activation. We find that, as previously shown, the CD56^bright^ NK cell subset is the most proliferative subset during activation with CIML stimulation and is also more sensitive to replication stress than CD56^dim^ NK cells. CD56^bright^ cells undergo caspase mediated apoptosis in response to replication stress induced by knockdown of MCM10 or treatment with a low dose of the replication inhibitor aphidicolin. The survival of T cells in response to CD3/CD28/IL-2 activation and replication stress remains relatively unaffected despite undergoing similar changes in replication fork speeds in response to aphidicolin treatment. We additionally found that perforin expression decreases with increasing replication stress, which provides a link between replicative stress and NK cell function. Finally, cell fate decisions due to DNA damage and replication stress are maintained in part by a balance of apoptosis and autophagy. We found that the CD56^dim^ subset undergoes autophagy in response to activation, though replication stress during activation does not further increase autophagy. Together, these results reveal insight into the mechanisms of NK cell deficiency produced by helicase deficiencies and replication stress and uncover novel aspects of NK cell biology and function.

## Results

### NK cell activation induces cell cycle dependent DNA damage

To better understand the effect of helicase deficiency on mature NK cells, we first sought to investigate proliferation and DNA damage in the context of activation. Activated NK cells were generated by overnight priming of healthy donor PBMCs with IL-15, IL-12, and IL-18 followed by low dose IL-2 for one to four days (CIML) (Fig. 1A) (11, 12, 15). Controls were given only low dose IL-2 or IL-15 for the same amount of time. Using BrdU, we found that CIML NK cells have a significantly increased frequency of cells found in S phase four days after stimulation relative to incubation with IL-15 only or no prime control (low dose IL-2 only) (Fig. 1B). Consistent with increased progression to S phase, we also confirmed that cell proliferation, a positive indicator for cellular activation (37), was increased in this CIML population when measured by CF-SE (Fig. 1C).

**Fig. 1.**
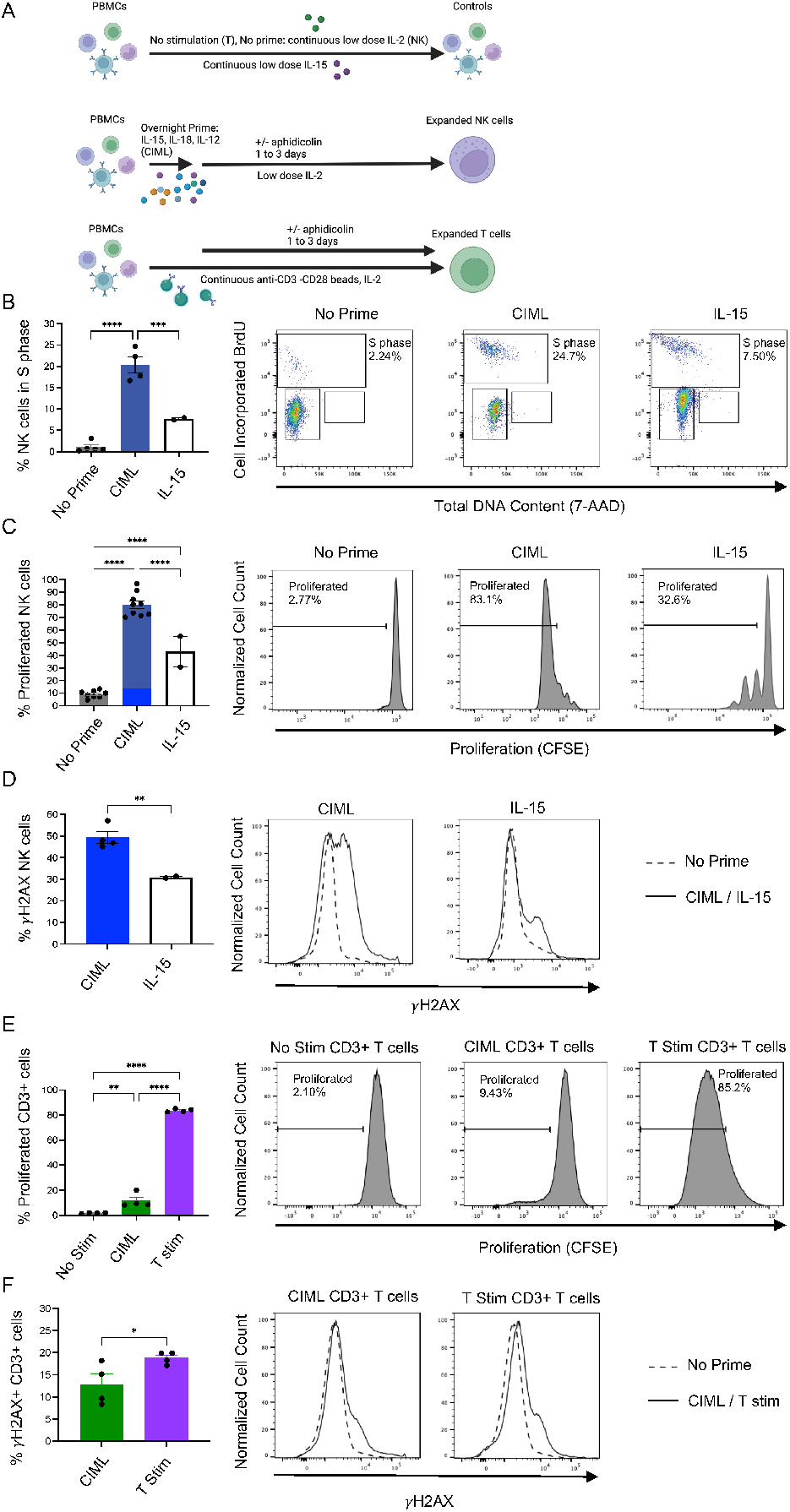
NK cell activation leads to cell proliferation, cell cycle progression, and DNA damage. A) Schematic of the PBMC stimulation protocol for negative control conditions (top), CIML NK cell activation (middle), and T cell activation (bottom). Negative controls received no stimulation for T cell experiments in Figures 1E-F, 20 U/mL IL-2 for NK cell experiments, or low dose (5 ng/mL) IL-15 for Figures 1B-D. CIML activated NK cells received IL-15, IL-12, and IL-18 overnight stimulation followed by low-dose IL-2 for 1-3 days in the presence or absence of aphidicolin. T cells received 30 U/mL IL-2 stimulation with anti-CD3 and anti-CD28 beads for 1-3 days in the presence or absence of aphidicolin. (B) Percentage of CD56+CD3– NK cells found in S phase by flow cytometric cell cycle analysis of BrdU and 7-AAD staining following activation with CIML cytokines or 5 ng/mL IL-15. n=4 biological replicates for no prime control, n=4 replicates for CIML conditions and n=2 replicates for IL-15 conditions, representative plot shown on right. (C) Percentage of proliferated NK cells based on CFSE staining by flow cytometry following activation with CIML cytokines or 5 ng/mL IL-15. n=2 (IL-15), 8 (no prime), and 9 (CIML) biological replicates; representative plot shown on right. (D) Percentage of *γ*H2AX CD56+CD3– NK cells by flow cytometry. Cells are gated based on unstained control. n=4 donors, representative plot shown on right. (E) Percentage of CD3+ T cells which have diluted CFSE dye in response to no stimulation, CIML activation, or T cell stimulation. (F) *γ*H2AX analysis by flow cytometry in CD3+ T cells activated with CIML cytokines or T cell stimulation. n=4 healthy donors were used in all T cell experiments. Cells are gated based on unstained controls. Data shown are mean*±*SEM.

To determine the effect of NK cell activation on conventional markers of DNA damage, we measured *γ*H2AX by flow cytometry following activation by CIML cytokines or IL-15 alone normalized to the no prime control. CIML stimulated NK cells had significantly more *γ*H2AX positive cells compared to IL-15 alone (Fig. 1D). To determine whether induction of DNA damage observed in cytokine primed NK cells is a conserved response amongst lymphocytes after activation, we compared activated NK cells with CD3+ T cells stimulated with anti-CD3/-CD28 magnetic beads and IL-2 (T stim). We also tested CIML conditions on T cells as an additional control to validate our T cell specific stimulation. CD4+ and CD8+ T cells consistently increased expression of activation markers CD69 and CD25 (38, 39) after three days of continuous T cell specific stimulation, whereas CIML cytokines did not have the same effect (Supp Fig 1A). We additionally confirmed using CF-SE staining that the proliferation of activated T cells was increased compared to no stimulation controls or CIML stimulation (Fig. 1E). Despite their robust activation and proliferation, T cells activated with T cell specific stimulation had only moderate increases in DNA damage measured by *γ*H2AX when normalized to an unstimulated control or compared to mild activation with CIML cytokines (Fig. 1F). In summary, stimulation of NK and T cells robustly induces proliferation in both populations to a similar extent when measured by CF-SE, however activation leads to a greater frequency of NK cells with DNA damage.

### Aphidicolin treatment impairs replication fork progression in NK cells and T cells

We sought to use mild replication stress to model the effect of helicase deficiencies on NK cells and T cells following their activation. At low doses such as 0.25 *μ*M and 0.5 *μ*M, aphidicolin induces replication stress, while higher doses like 1 *μ*M and 5 *μ*M result in cell cycle arrest (40, 41). Following CIML cytokine priming overnight, NK cells were cultured with low dose IL-2 in the presence or absence of 0.25 *μ*M aphidicolin for 72 hours. NK cell survival in the absence of activation was tested us-ing condition with no priming but continuous low dose IL-2. Induction of replication stress with aphidicolin leads to a similar phenotype as previously reported in cells from helicase-deficient patients, particularly the MCM10-deficient individual (23), including DNA damage and cell cycle arrest in early S phase (Supp. Fig. 2A). In parallel, we performed T cell stimulation as previously described and including a continuous low dose IL-2 control, followed by aphidicolin treatment for the same length of time as NK cells.

Activation of NK cells without aphidicolin did not induce cell death as detected by propidium iodide (PI) staining. However, low dose replication stress during activation significantly increased the frequency of PI+ NK cells (Fig. 2A). In contrast, while T cell activation increased the frequency of PI+ T cells relative to the low dose IL-2 control, there was no significant increase in this population with 0.25 *μ*M aphidicolin (Fig. 2A). We sought to determine whether the differential response we observed between activated T cells and NK cells was due to differences in the effect of aphidicolin on replication fork speed. DNA fiber assays were performed on activated T cells or NK cells in the presence or absence of 0.25 *μ*M aphidicolin. Replication fork speed in the absence of aphidicolin was not significantly different between NK cells and T cells. Notably, two hours of aphidicolin treatment in the presence of cell type specific activation similarly decreased fork speeds in both NK and T cells (Fig. 2B), suggesting that the differences in response to 0.25 *μ*M aphidicolin were not due to baseline differences in fork speed between NK cells and T cells and not due to differential effects of aphidicolin on DNA replication fork progression.

**Fig. 2.**
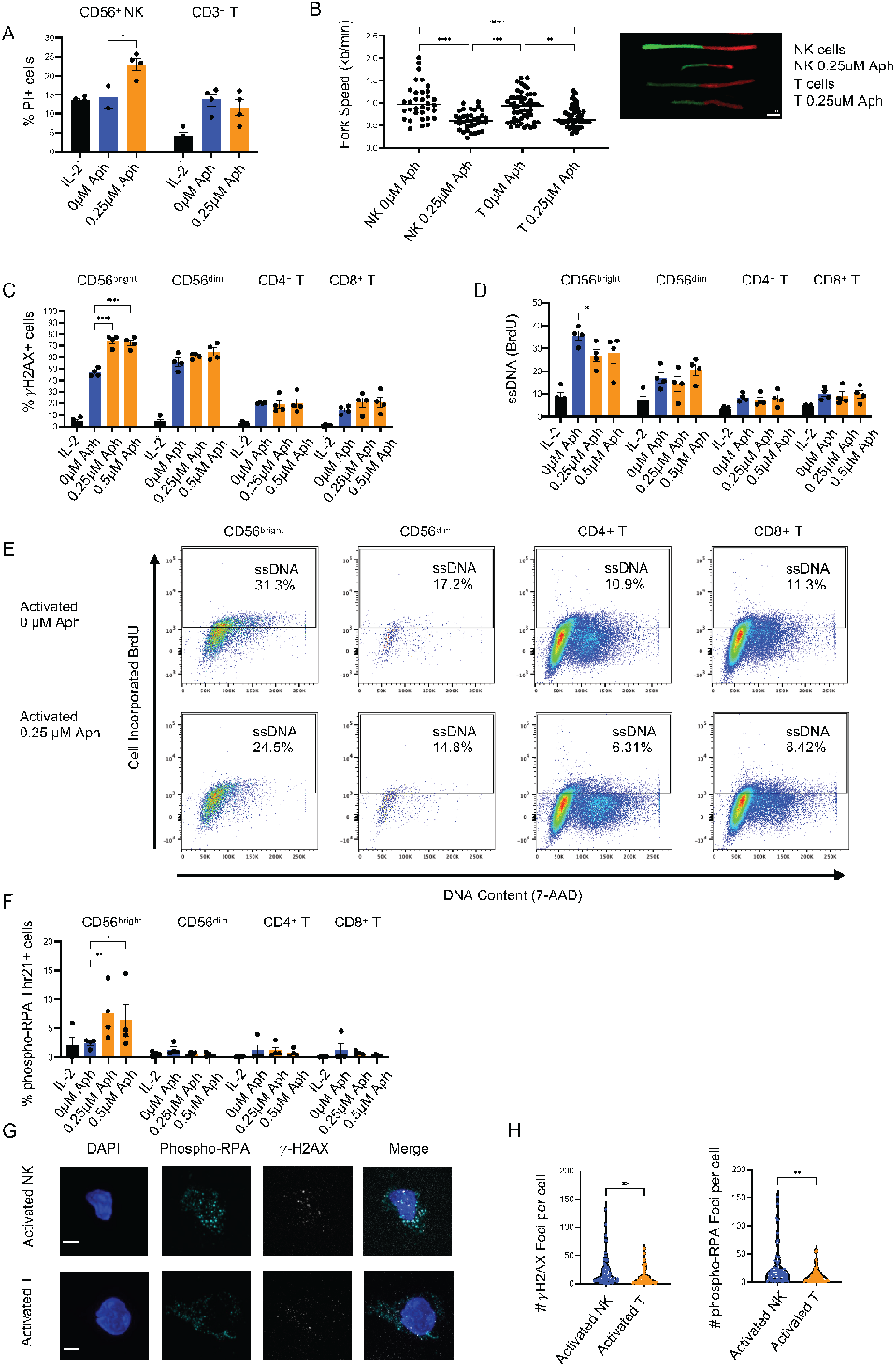
Aphidicolin treatment impairs replication fork progression in NK cells and T cells. A) Percent propidium iodide (PI) positive CD56+CD3– NK cells and CD3+ T cells following low dose IL-2 or NK (CIML) or T cell activation with or without aphidicolin treatment. (B) Fork speed in kilobases per minute of enriched stimulated NK cells or T cells from a DNA fiber analysis assay. Cells were incubated for two hours with 0.25 *μ*M aphidicolin following overnight cytokine stimulation for enriched NK cells or continuous anti-CD3 -CD28 beads with 30 U/mL IL-2 for T cells. Cells from two healthy donors were pooled to make the DNA fiber slides. (C) Percentage of *γ*H2AX+ cells in NK and T cell subsets with or without low dose aphidicolin following activation with low dose IL-2 or CIML activation (NK cells) or T cell activation. (D) Percentage of BrdU+ cells without DNase treatment (exposed ssDNA) measured by flow cytometry. (E) Example flow plots are shown with BrdU+ gating strategy based on BrdU FMO control with DNA content on the X-axis. (F) Phospho-RPA (Thr21)+ cells in NK and T cell subsets with or without low dose aphidicolin following activation with low dose IL-2 or CIML activation (NK cells) or T cell activation. Cells are gated based on fluorescence minus one or no stain controls. (G) Representative confocal microscopy images from CIML activated NK cells (top) and activated T cells (bottom). (H) Quantification of *γ*H2AX or pRPA (Thr21) foci in CIML NK cells and activated T cells. n=4 biological replicates. Two-way ANOVA comparing cell subsets was performed for multiple comparisons. Data shown are mean*±*SEM.

We predicted that the addition of aphidicolin would further increase the presence of *γ*H2AX induced by activation of NK cells. As CD56^bright^ and CD56^dim^ NK cells have differential proliferative responses to CIML activation (42), we measured these subsets, and CD4+ and CD8+ T cell subsets, independently following cell type specific activation or low dose IL-2.*γ*H2AX intensity was significantly increased in the CD56^bright^ subset in response to aphidicolin (Fig. 2C). The CD56^dim^ population had increased DNA damage in response to activation but did not further accumulate damage with the addition of replication stress (Fig. 2C). Similarly, both CD4+ and CD8+ T cell populations accumulated low amounts of DNA damage with stimulation and maintained this low level even with replication stress (Fig. 2C). These data demonstrate that CD56^bright^ NK cells are specifically sensitive to replication stress.

We sought to measure additional indicators of replication stress and DNA damage. Exposed single stranded DNA (ss-DNA) is present during homology directed repair and in the process of proliferation and replication activities. The replication protein A (RPA) binding stabilizes ssDNA and facilitates downstream DNA damage repair activities (43). To measure the presence of ssDNA, we performed flow cytometry BrdU assays in the absence of the commonly used DNA denaturing step. In CD56^bright^ NK cells, we found an increase in exposed ssDNA in response to activation (Fig. 2D, E). The addition of low dose aphidicolin decreased the percentage of BrdU positive cells, likely due to decreased cell cycle progression (Fig. 2D, E). CD56^dim^ cells also increased ssDNA in response to activation, though not to the same degree as the CD56^bright^ subset. Both subsets of T cells maintained low levels of ssDNA that were seemingly independent of activation or replication stress (Fig. 2D, E). We found a trend similar to *γ*H2AX distribution when we analyzed phosphorylated RPA (phospho-RPA Thr21) by flow cytometry and found that the CD56^bright^ NK cell subset had increased phospho-RPA (Thr21) following treatment with 0.25 *μ*M aphidicolin (Fig. 2F). In contrast, the CD56^dim^ subset and T cell subsets maintained low levels of phospho-RPA regardless of activation or replication stress (Fig 2F). Confocal microscopy showed a greater number of *γ*H2AX and phospho-RPA (Thr21) foci in activated NK cells than T cells (Fig. 2G) and this observation was supported by quantification (Fig. 2H). These data indicate that the CD56^bright^ subset is affected by mild replication stress while undergoing CIML activation to a greater extent than CD56^dim^ NK cells or T cell subsets being activated by T cell specific stimulation. They further show that, despite equivalent replication fork speeds and response to aphidicolin, NK cells and T cells respond differently to mild replication stress.

### Replication stress leads to apoptosis of activated NK cells

Next, we sought to further investigate the mechanisms of increased NK cell death in response to activation and replication stress. The tumor suppressor transcription factor p53 plays a role in controlling cell fate, including cell cycle arrest, repair, senescence, or apoptosis (44). We investigated phospho-p53 (Ser15) and p21, which is a downstream target of phospho-p53, by flow cytometry for NK and T cell subsets. As shown in Fig. 3A, the CD56^bright^ NK cells had highest expression of phospho-p53 following activation, with or without replication stress. The CD56^dim^ NK cell subset had moderate levels of phospho-p53 that were increased relative to no prime controls. However, T cells maintained low levels of phospho-p53 and these were independent of aphidicolin treatment, and neither T cells nor NK cells had significant phospho-p53 expression in response to low dose IL-2 controls. Analysis of mean fluorescent intensity (MFI) displayed a similar trend, where activated CD56^bright^ cells have increased p21 compared to no prime controls, without further impacts by replication stress (Supp. Fig 3A). p21 MFI in CD56^dim^ NK cells and CD3+ T cells was unchanged by activation or replication stress (Supp. Fig 3A). Next, we measured pan-caspase expression by flow cytometry to determine signaling pathways resulting in apoptosis. NK cells had greater caspase activation with increasing replication stress, whereas T cells maintained a low level of caspase activity even in the presence of aphidicolin and both cell types had low caspase expression in low dose IL-2 conditions (Fig. 3B). This result was further confirmed by MitoNIR staining, which decreases in intensity with loss of mitochondrial membrane potential, indicating cell death (45). Here, we found moderate increases in the percentage of cells that were MitoNIRlow with replication stress for both CD56^bright^ and CD56^dim^ populations, with both populations having higher frequencies of MitoNIRlow cells than either T cell subset (Fig. 3C). CD4+ and CD8+ T cells did not have increased MitoNIRlow populations following aphidicolin treatment.

**Fig. 3.**
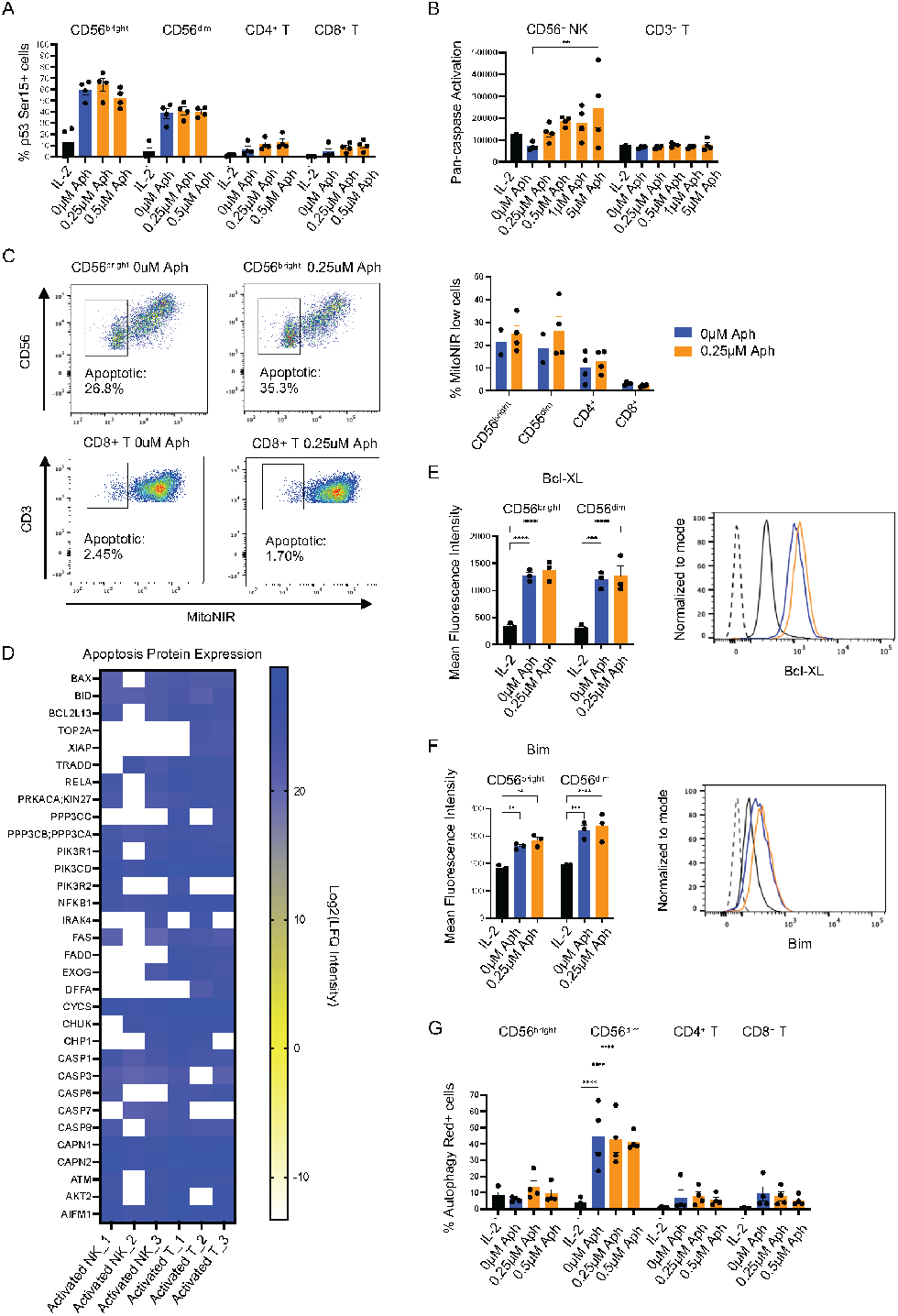
Replication stress leads to apoptosis of activated NK cells. A) Percent p53 phospho-Ser15 positive cells in NK and T cell subsets with or without low dose aphidicolin following activation with low dose IL-2 or CIML activation (NK cells) or T cell activation. No statistical significance was calculated within each subset compared to stimulation control. n=4 biological replicates. (B) Fluorescence intensity of pan-caspase dye was collected for CD56+ NK cells and CD3+ T cells. n=4 biological replicates. (C) Percent MitoNIRlow cells for NK and T cell subsets with and without aphidicolin. Gating strategy was based on IL-2 only control. n=4 biological replicates. (D) Proteomic analysis of the Log2(LFQ intensity) of selected pro-apoptotic proteins from isolated NK cell and T cells. n=3 biological replicates. (E) and (F) Flow cytometry of pro-apoptotic markers was conducted on n=3 healthy donors. (G) Percent Autophagy Red positive cells were calculated for each NK and T cell subset. Two-way ANOVA comparing cell subsets was performed for multiple comparisons.

We also conducted proteomics analysis of activated NK and T cells and found variable differences in pro-apoptotic proteins levels (Fig. 3D). Notably, NK cells upregulated CASP7 and CASP3 compared to T cells, however, several proteins were not detected in activated NK cells, such as TOP2A, XIAP, FADD, and DFFA (Fig. 3D). To complement the proteomics analysis, we investigated anti-and pro-apoptotic proteins by flow cytometry in activated NK cells with and without replication stress, including Bcl-XL (Fig. 3E) and BIM (Fig. 3F), respectively. Here, we find that both subsets of NK cells had increased protein expression with CIML activation relative to low dose IL-2 only, while the addition of aphidicolin to induce replication stress did not further increase protein expression. Thus, NK cell activation, and not just the addition of replication stress, induces the upregulation of both anti- and pro-apoptotic factors, indicating a readiness for apoptosis to occur.

Finally, we asked if the process of autophagy could be playing a role in lymphocyte response to replication stress. Autophagy is an important recycling process necessary for maintaining cellular homeostasis (46). Autophagy may be activated by the same stresses as apoptosis, such as in response to replication stress and DNA damage (46, 47). Here, we used the fluorescent probe Autophagy Red, which is detected in the lipid membrane of autophagosomes and autolysosomes. We found that the CD56^dim^ subset had the highest percentage of cells undergoing autophagy following activation in the presence or absence of replication stress (Fig. 3G). Relative to CD56^dim^ NK cells, CD56^bright^ NK, CD4+ and CD8+ T cells maintained low levels of autophagy following activation with or without replication stress (Fig 3G). Through phospho-p53 signaling, loss of mitochondrial membrane potential, and caspase activation, apoptosis occurs in CD56^bright^ NK cells in response to activation. Caspase activity was particularly sensitive to replication stress in NK cells. The CD56^dim^ subset of NK cells undergoes autophagy rather than apoptosis in response to activation and replication stress. Together, these data indicate that NK cells, especially the CD56^bright^ subset, exhibit differential sensitivity to replication stress compared to T cells.

### Differential expression of CMG complex members in NK and T cells

The function of CMG helicase proteins is tightly linked to their expression (31–33), and loss-of-function variants in MCM4, GINS1, MCM10 and GINS4 lead to immune deficiency. We hypothesized that differences in the cell-intrinsic expression of CMG helicase complex proteins could account for the differential response to replication stress between NK cells and T cells. We performed Western blotting on isolated resting NK cells and T cells from three healthy donors. GINS4 and CDC45 were selected as representative proteins within the CMG helicase. We found no difference in the relative expression of GINS4 (Fig. 4A) and CDC45 (Fig. 4B) in resting NK and T cells from three donors when normalized. Next, we activated NK cells with high and lose doses of IL-15 to induce activation with less cell death than CIML conditions (10 ng/mL and 1 ng/mL, respectively) and T cells with soluble anti-CD3 and -CD28 with soluble IL-2. Activation led to increased expression of GINS4 with a greater increase in expression in T cells than NK cells (Fig. 4C). In contrast, we did not detect a consistently significant increase in CDC45 protein expression between conditions due to variability in expression between donors (Fig. 4D). To perform further unbiased comparison of changes in protein expression, activated NK and T cells from three healthy donors were evaluated by proteomics analysis after CIML or T cell activation, respectively. Here, we consistently found significantly increased MCM proteins in activated T cells compared to NK cells (Fig. 4E, F, G). Similarly, we found increased levels of other canonical proteins required for replication, including PCNA and POLD1, in T cells but not NK cells (Fig. 4H, I). We included ACTB as a housekeeping protein control, which did not identify significant differences between activated NK and T cells (Fig 4J).

**Fig. 4.**
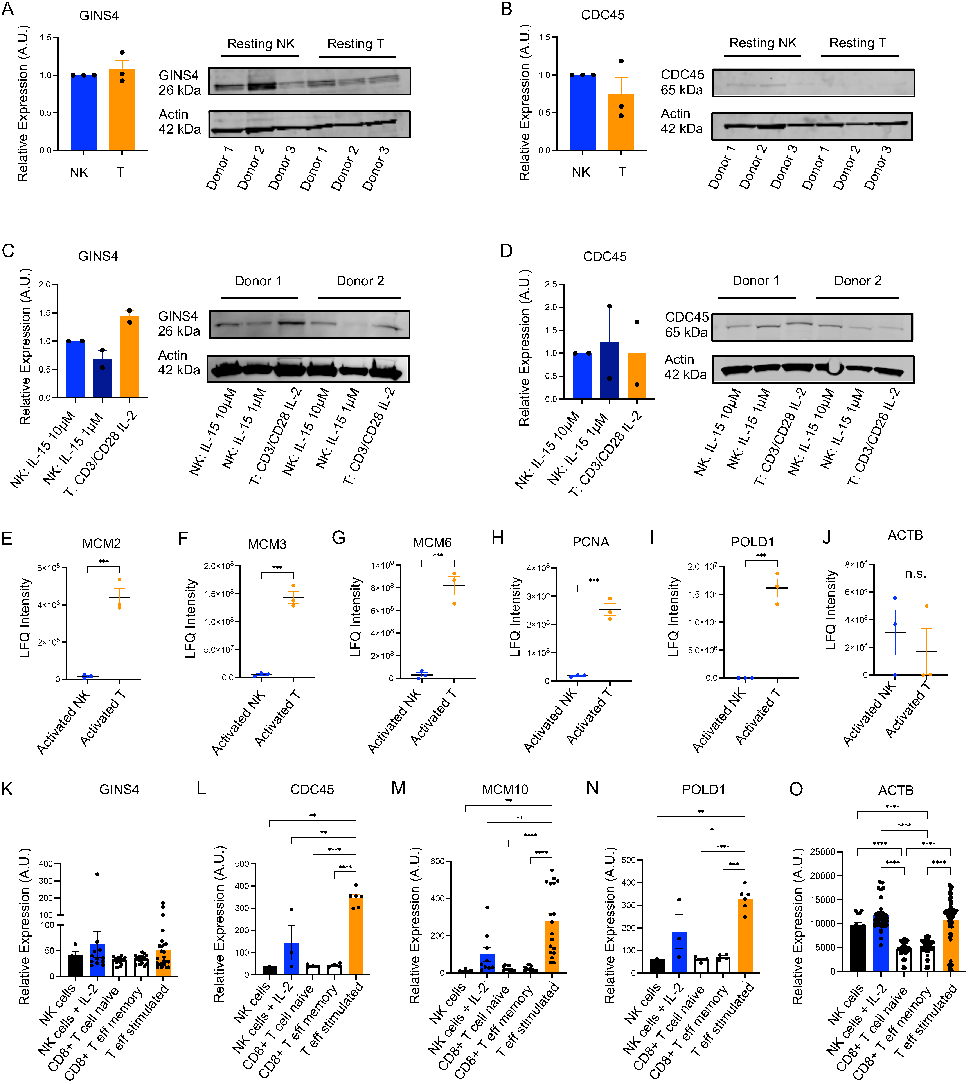
Differential expression of CMG complex members in NK and T cells. Western blot of resting T cells or NK cells from 3 healthy donors blotted for GINS4, CDC45 and actin (for normalization). Protein was normalized first to actin for each lane then normalized to NK cells. No statistical significance was found between NK and T cells. (C) and (D) Western blotting of activated NK cells (IL-15) and T cells (CD3/CD28 IL-2) from two healthy donors was conducted using the same antibodies and analysis as resting samples. No statistical analysis was conducted as only two healthy donors were included. (E-I) Proteomic LFQ intensity data was plotted for selected MCM proteins and canonical cell cycle proteins. One-tailed T tests were conducted to determine statistical significance for each protein. (J-M) Relative expression in arbitrary units for RNA abundance of given helicase proteins and replication genes was collected and analyzed from BioGPS. One-way ANOVA was used to test for differences.

Next, we analyzed the primary cell RNA-seq atlas data from BioGPS (48), as it allowed us to compare helicase and DNA replication factors expressions in resting versus IL-2 activated NK cells and naïve versus stimulated T cells. We found a consistent pattern of increased gene expression in response to T or NK cell activation in *GINS4* (Fig. 4K), *CDC45* (Fig. 4L), *MCM10* (Fig. 4M), and *POLD1* (Fig. 4N). *ACTB*, included as a control protein, shows similar increases for stimulated NK and T cells (Fig 4O). Notably, stimulated T cells tended to have increased gene expression in response to activation to a greater extent than NK cells, a trend that was consistent for other helicase-related genes including *GINS1, GINS3, MCM2, MCM6, MCM7*, and *POLA1* (Supp. Fig. 4A-F). Thus, differential gene expression changes, namely increased expression in response to activation, could contribute to the greater ability of T cells to withstand mild replication stress relative to NK cells.

### Replication stress impairs the cytotoxic capacity of primary human NK cells

Replication stress induced apoptosis and autophagy in CD56^bright^ and CD56^dim^ cells respectively (Fig. 3), though some cells do survive after presumably repairing DNA damage and restarting the cell cycle. We investigated the cytotoxic capacity of NK cells that had survived replication stress by measuring lytic effector molecules and directly testing cytolytic function of primary NK cells against target cells. We found that activation in the presence of aphidicolin led to a decreased frequency of granzyme B positive CD56^dim^ and CD56^bright^ cells in a dose-dependent manner (Fig. 5A). CD56^dim^ cells followed this trend for granzyme A, which triggers cell death through mitochondrial breakdown (49), however the CD56^bright^ subset were less affected by replication stress when considering the percentage of granzyme A positive cells (Fig. 5B). The percent of perforin positive CD56^dim^ and CD56^bright^ NK cells also decreased with increasing replication stress (Fig. 5C). Interestingly, IFN-*γ* production of CD56^bright^ NK cells was unaffected by replication stress and even 5 *μ*M aphidicolin for three days did not affect IFN-*γ* production by surviving cells, despite CD56^bright^ NK cells being the subset most strongly affected by replication stress (Fig. 5D). Finally, we directly tested the lytic capacity of replication stress-affected NK cells on K562 leukemic target cells. Using healthy donor activated and enriched NK cells from PBMCs, we found decreased percent specific lysis against target cells, though no statistical significance was determined (Fig. 5E). Thus, in addition to the replication stress in activated NK cells resulting in apoptosis and autophagy, the surviving cells are also mildly affected showing decreased expression of effector proteins that can lead to decreased cytotoxicity against target cells.

**Fig. 5.**
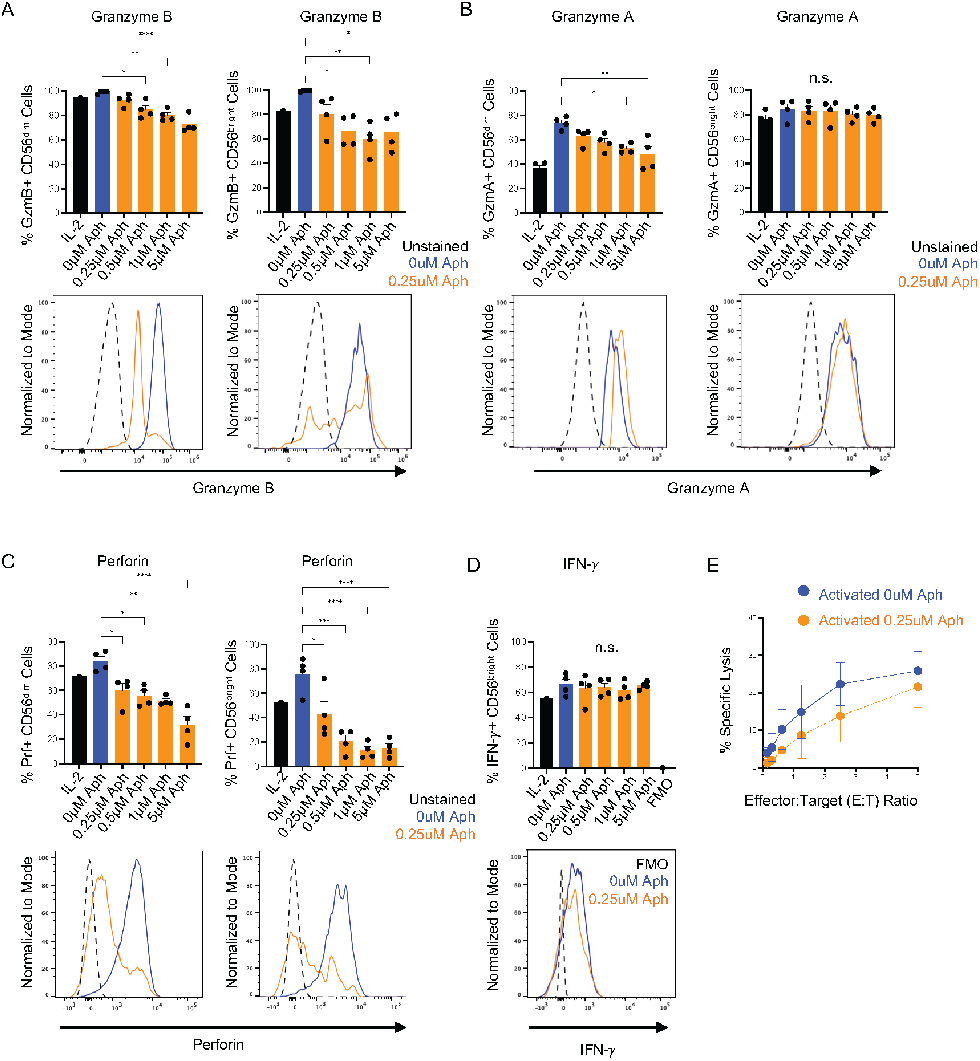
Replication stress impairs the cytotoxic capacity of primary human NK cells. Four healthy donors were used to analyze intracellular granzyme B and granzyme A respectively and gated for CD56dim and CD56^bright^ cells. One-way ANOVA calculated statistical significance between stimulated control and those with aphidicolin. Results of significance compared to IL-2 only are not shown. Below bar graphs are example histograms featuring unstained control, stimulated control, and 0.25 *μ*M aphidicolin conditions. (C) Intracellular perforin was investigated for four healthy donors and gated on CD56dim and CD56^bright^ cells. Representative histograms include unstained and stimulated controls and the lowest dose of 0.25 *μ*M aphidicolin condition. One-way ANOVA calculated statistical significance, but results compared with IL-2 only condition is not shown. (D) Cytokine production for IFN-*γ* was conducted for CD56^bright^ cells, including a fluorescence minus one negative control. No statistical differences were calculated by One-Way ANOVA. (E) 51Cr release assay to evaluate the cytotoxic function against K562 cells for five healthy donors. Cells were enriched using the StemCell kit described in the Methods section. Data shown are mean*±*SEM of technical replicates. Data are representative of three independent experimental repeats.

### Helicase protein knockdown leads to increased NK cell apoptosis

Previously, our lab tested the effect of MCM10 knockdown in the NK92 NK cell line using CRISPR-Cas9 (23). MCM10-knockdown (KD) NK92 cells exhibited increased frequency of cells in early S phase and increased doubling time, indicative of replication stress (23). To further model the MCM10 loss-of-function variants in an NK cell line with a terminally mature NK cell phenotype (50), we targeted MCM10 expression using shRNA in the YTS human NK cell line. Western blot confirmed decreased protein expression by about 80% in the MCM10 knockdown (MCM10-KD) cell line compared to scramble shRNA control (Fig. 6A). We also found a higher proportion of knockdown cells were PI positive by flow cytometry, indicating poor cell survival of MCM10-KD cells (Fig. 6B). BrdU incorporation to investigate cell cycle revealed a significantly higher percentage of MCM10-KD YTS cells in S phase, specifically early S phase (Fig. 6C). Taken together, these data recapitulate previously described impairment in cell cycle progression with a greater number of cells in early S phase resulting from loss of MCM10 function (23) and further revealed impaired cell survival resulting from MCM10-knockdown.

**Fig. 6.**
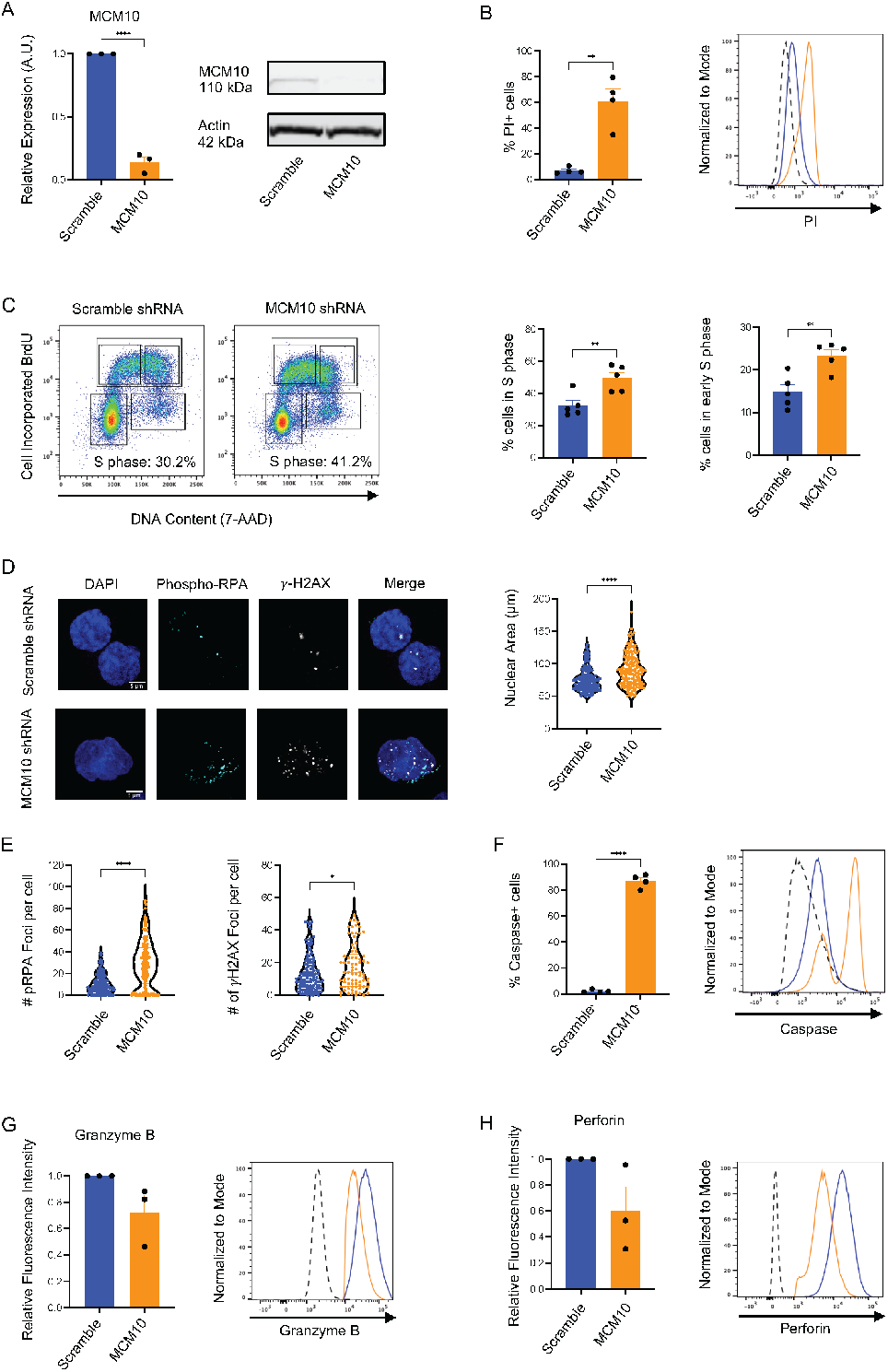
MCM10 knockdown leads to increased NK cell apoptosis. Validation of MCM10 knockdown was conducted by Western Blotting using anti-MCM10 and anti-actin antibodies. n=3 technical replicates. (B) Relative percent propidium iodide positive cells was analyzed by flow cytometry using fluorescence minus one (FMO) controls. Representative histograms are shown. n=4 technical replicates. (C) Representative cell cycle graphs with gating strategy based on scramble shRNA and quantification of the frequency of cells in S phase or early S phase. n=5 technical repeats. (D) Representative images from confocal microscopy of MCM10-KD or scramble (control) shRNA YTS cells with DAPI staining and antibodies against phospho-RPA and *γ*H2AX. Nuclear area quantification of knockdown cells compared to controls. (E) Number of foci of phospho-RPA and *γ*H2AX compared to scramble shRNA controls. (F) Relative percent caspase positive cells using FMO controls. n=3 technical replicates. Student’s T test was used to compare knockdown with scramble shRNA control. (G) and (H) Intracellular flow cytometry for granzyme B and perforin. n=3 technical replicates. Representative histograms are shown. No statistical significance was detected between MCM10-KD and controls. Scramble, scramble shRNA YTS cells; MCM10, MCM10-KD YTS cells.

Next, we sought to quantify the effect of MCM10 knockdown on markers of replication stress. We performed confocal imaging of nuclear area, phospho-RPA (Thr21) foci, and *γ*H2AX foci in MCM10-KD and scramble control lines. Image analysis revealed increased nuclear area (Fig 6D), phospho-RPA foci, and *γ*H2AX foci (Fig. 6E). This is in line with the MCM10-deficient patient where increased nuclear area and *γ*H2AX foci were seen (23). We also measured apoptosis in the MCM-KD and control cells and found the percentage of pan-caspase positive cells was increased in MCM10-KD (Fig. 6F). Given the impaired cytotoxic function of primary NK cells activated and treated with aphidicolin, we sought to determine the effect of MCM10-KD on YTS cell function. There was a consistent decrease in intra-cellular granzyme B (Fig. 6G) and perforin (Fig. 6H) relative to scramble shRNA controls, however this decrease was not statistically significant (Fig. 6G, H). Together, these data indicate that decreased expression of helicase proteins can recapitulate the phenotype described earlier in replication stress-induced activated healthy donor NK cells. Thus, helicase deficiency as a result of loss of function or decreased expression of core replisome proteins results in replication stress, which is ultimately the driver of apoptosis and decreased cytotoxicity in NK cells but not in activated T cells.

## Discussion

The association between partial loss-of-function variants in genes encoding the CMG helicase and NKD have been well documented (19–23). Cellular phenotypes of these variants include cell cycle arrest, DNA damage, and replication stress (19–23). While these variants impair NK cell development, they also negatively affect the proliferative response of mature NK cell subsets (19, 20, 22, 23), while T cell responses are often intact or less affected than NK cells (19, 20, 23). Here, we use DNA polymerase inhibition with aphidicolin to increase replication stress in primary NK cells and gene knockdown to recapitulate the effects of helicase deficiency in an NK cell line. Aphidicolin was used throughout our study as a means to delay replication by inhibiting DNA polymerase. Our lab and other have shown that limitations to helicase function through helicase deficiency leads to DNA damage, replication stress and cell cycle arrest, which is a phenotype that also appears when aphidicolin is used at low doses. We show that replication stress induced by aphidicolin strongly impacts the CD56^bright^ NK cell subset by increasing *γ*H2AX and phosphorylated RPA and ultimately triggers an apoptotic program in NK cells but not T cells. These data provide mechanistic insight into the nature of immunodeficiency in individuals with helicase variants. They also provide interesting new biological insight into how NK cells and T cells differ in response to replication stress as well as helicase protein expression, despite both cell types being cytolytic effector cells.

We explored helicase protein expression in resting and activated NK and T cells and found that T cells upregulated helicase genes and proteins to a higher degree than NK cells. Increased CMG helicase protein expression has been associated with increased proliferation, especially in relation to cancer and reviewed in full elsewhere (51–56). Here, we show that activated NK cells and T cells proliferate at a similar rate and have similar replication fork speeds in our in vitro assays, yet T cells have a greater increase in helicase protein expression in response to activation. Perhaps T cell control of helicase proteins during proliferation offers protection against DNA damage and replication stress by allowing multiple dormant origins to fire at once and maintaining adequate DNA damage response signaling (57).The MCM paradox states that excess inactive MCM proteins are loaded onto chromatin, yet only a fraction of these are licensed into active replication forks during S phase (58). A leading theory towards understanding this paradox hypothesizes that the excess MCM proteins are necessary to trigger dormant origins during replication stress (58). Thus, T cell upregulation of helicase proteins during activation could combat replication stress by initiating dormant replication origins. Conversely, NK cells could use a lower threshold of helicase proteins during proliferation but are left at risk of programmed cell death in response to stimulatory signals.

It is pertinent to note that activated NK cells have significant DNA damage when compared to unstimulated controls, indicating that NK cells during viral infection will likely accumulate DNA damage just by being activated and could undergo activation-induced cell death (AICD). In support of this, Terren et al demonstrated that CIML activation of NK cells resulted in decreased viability, likely due in part to failed mitochondrial dynamics and mitophagy. Apoptosis resulting from activation was further confirmed with accumulation of superoxide and release of cytochrome c in NK cells (59). T cells, particularly CD8+ T cells, undergo AICD as a form of peripheral immune regulation with signaling through Fas/FasL or TNFR2, which is particularly important for the prevention of autoimmune disease (60, 61). However, a threshold for AICD in NK cells after activation may be contributing to our results where we find a higher sensitivity to DNA damage and replication stress. Research to clearly define AICD in NK cells would further this hypothesis and improve our biological understanding of NK cells responses during human disease.

Understanding natural killer cell requirements for replication, especially under conditions of stress, is paramount for deciphering the role of the CMG helicase in these cells. Our work and previous helicase NKD studies have determined that NK cell maturation is blocked, particularly through apoptosis of the CD56^bright^ population. Gineau et al found that CD56^bright^ NK cells from MCM4 deficient individuals had chromosomal aberrations, indicative of replica-tion stress, and had increased apoptosis (22). In our data, aphidicolin-induced replication stress resulted in apoptosis of the CD56^bright^ population, which left behind a small population of cells that can survive replication stress yet have impaired cytotoxicity, as demonstrated in Figure 5. Considering our results here, we have identified a connection between helicase deficiency and replication stress leading to NK cell-specific dysfunction. Our work shows that human NK cells and T cells have differential sensitivities to replication stress, and additional research should further study the connection between helicase proteins and disease. Better understanding of how cytokine-induced activation affects NK cell survival, and the effect of mild replication stress on activated NK cells, can help better design strategies for the use of immunotherapeutic NK cells.

## Methods

### Cell isolation and cell lines

PBMCs were isolated by density gradient centrifugation from the buffy coat of healthy donors from the New York Blood Bank using Ficoll-Paque (Fisher Sci Cat# 45-001-750). PBMCs were stored frozen in 90% FBS 10% DMSO and thawed for each experiment. Once thawed, PBMCs were resuspended and maintained in RPMI 1640 medium (Thermo Cat# 11875135) supplemented with 10% heat-inactivated human AB serum (GeminiBio Cat# 100-512), 1% penicillin-streptomycin (PenStrep; Thermo Cat# 15140163), 2mM Glutamax (Thermo Cat# 35050079), 1X non-essential amino acids solution (NEAA) (Thermo Cat# 11140076), 1mM sodium pyruvate (Thermo Cat# 11360070), and 1M HEPES (Thermo Cat# 15630130). YTS cell lines, which are derived from malignant NK cell leukemia/lymphoma (38), were maintained with the same media as PBMCs, except with 10% FBS (Atlanta Biologicals Cat# S11150) instead of human serum. K562 cells were maintained in RPMI medium supplemented with 10% FBS, 1% PenStrep, and 2mM GlutaMAX at 37C and 5% CO2. All cell lines were confirmed negative for mycoplasma.

### NK cell and T cell stimulation

Frozen PBMCs were thawed and briefly rested in human serum-supplemented RPMI medium. To generate activated NK cells, cytokines were added to the media (IL-15 100 ng/mL, IL-18 50 ng/mL, IL-12 10 ng/mL) for overnight stimulation. PBMCs were then washed three times and resuspended with fresh media and low dose IL-2 (20 U/mL) for expansion phase in a round-bottom 96-well or 24-well plate. Aphidicolin (0.25 *μ*M, 0.5 *μ*M, 1 *μ*M, or 5 *μ*M) (Sigma Cat# A4487) was also added at this time. To generate activated T cells from PBMCs, fresh media was supplemented with 30 U/mL of IL-2 and a 1:1 ratio of anti-CD3 and -CD28 Dynabeads (Thermo Fisher Cat# 11131D) and similarly treated with aphidicolin. Cells were harvested one or three days after aphidicolin and used for flow cytometry with surface markers to identify NK and T subsets or enriched using StemCell Negative Isolation kits (EasySep Human NK Cell Isolation Kit StemCell Cat# 17955 and EasySep Human T Cell Isolation Kit StemCell Cat# 17951).

### Flow cytometry

Phenotypic flow analyses were conducted on both PBMCs and cell lines using multicolor flow cytometry. Cells were washed in PBS then surface stained with fluorochrome-conjugated antibodies and viability dye at 4C for 20 minutes while protected from light. Cells were washed again then fixed and permeabilized for intracellular staining using the FoxP3 nuclear staining kit (Fisher Scientific Cat# 00-5523-00) according to manufacturer’s instructions. Fluorescence minus one and unstained controls were included for all experiments. To measure IFN-g, 10 ug/mL Brefeldin A (Biolegend Cat# 420601) was added during stimulation to inhibit protein transport. Data were acquired on a BD Fortessa and data analyzed with Flow Jo v.10.9.0 (BD Biosciences).

### Cell cycle analysis

Cell cycle analyses were conducted with the BrdU assay kit from BD Biosciences according to manufacturer’s instructions. Cells were incubated with 10 *μ*M BrdU for two hours followed by extracellular marker staining, fixation, denatured with DNase, and incubation with anti-BrdU (BD Biosciences Cat# 559619) and any other intracellular markers. Finally, cells were stained with 7-AAD for five minutes, then analyzed on a BD Fortessa cytometer. Results were analyzed with Flow Jo and the frequency of cells found in cell cycle phases (G1/G0, S, G2/M) was determined by gating BrdU-7-AAD intensity.

### Confocal microscopy

Cells were briefly adhered to #1.5 imaging chambers followed by fixation and permeabilization using Cytofix/Cytoperm (BD Biosciences Cat# 554714). Immunostaining was performed with phospho-RPA (Thermo Cat# PA5-104809) followed by goat anti-rabbit Alexa Fluor 488 (Thermo Cat# A32731), then γH2AX Alexa Fluor 647 (BD Biosciences Cat# 560447). DAPI staining was performed for 10 minutes prior to imaging. Images were acquired with a 100X 1.46 NA objective on a Zeiss AxioObserver Z1 microscope stand equipped with a Yokogawa W1 spinning disk by imaging cell volumes with a 0.17 *μ*m Z step size. Illumination was by 405 nm, 488 nm, and 647 nm solid state lasers and detection was by a Prime 95B sCMOS camera. Data were acquired in SlideBook software (Version 6, Intelligent Imaging Innovations) and exported as OME-TIFF files for further analysis. Images were analyzed as Z projections in Fiji (39) using the “Analyze Particles” plug-in with a minimum size of 0.05 *μ*m^2^. Data were exported to Prism 9 (GraphPad Software) for graphing.

### shRNA Lentiviral Transduction

YTS cells were maintained in culture medium in growth phase and 10^6^ cells were plated in 6-well plate 24 hours prior to shRNA viral transduction. Cells were washed with sterile PBS and resuspended in 400 uL fresh complete media with 100 L TransDux MAX Lentivirus Transduction Reagent (SystemBio Cat# LV860A-1). Viral supernatant for non-targeting (SMARTvector Lentivirus shRNA hEF1a TurboGFP, Horizon Discovery Cat# S02-005000-01) and MCM10 (SMARTvector Lentivirus shRNA MCM10 TurboGFP, Horizon Discovery Cat# V3SH7669) were added to respective wells and the plate was spun at 1500g for 90 minutes at 35C. Cells were then incubated for 48 hours at 37C. Antibiotic resistance selection with puromycin began 72 hours after transduction starting at 0.5 ug/mL and gradually increasing to 2 ug/mL over the course of two weeks. Cells were GFP+ sorted after 2 ug/mL puromycin selection. MCM10 protein knockdown was validated by Western blot.

### DNA fiber analysis

DNA fiber analysis was carried out as previously described (62). Briefly, cells were incubated with 25 M IdU for 30min, washed 3 times with warm PBS and incubated with 25 *μ*M CIdU for another 30min. Fibers were stretched on slides and stained with CldU (Bio-rad Cat# OBT0030), and IdU (BD Cat# 347580) antibodies. Slides were mounted with ProLong Gold Antifade mounting medium and dried overnight. The fiber tracks were imaged on a Nikon Eclipse 90i microscope fitted with a PL Apo 40X/0.95 NA objective and measured using ImageJ software.

### Proteomics analysis

NK cells and T cells were stimulated using IL-15, IL-12, and IL-18 or anti-CD3/CD28 beads with IL-2 respectively and harvested three days after stimulation. Magnetic Dynabeads were removed from T cell samples using the DynaMag-2 Magnet (Invitrogen Cat# 12321D) and then NK and T populations were enriched using StemCell Negative Isolation kits. Cells were briefly washed with cold PBS twice and transferred to 1.5mL eppendorf tubes. Samples were flash frozen using liquid nitrogen and delivered to the Proteomics and Macromolecular Crystallography core at the Columbia University Herbert Irving Comprehensive Cancer Center (HICCC). Label-free protein quantification using LC-MS was conducted on the samples. Data was analyzed using Microsoft Excel and graphs were made with Prism 10.0.0 (GraphPad).

### NK cell cytotoxicity assays

Cytotoxicity of enriched activated NK cells from PBMCs from five healthy donors against K562 erythroleukemia target cell line was performed by four-hour Cr51 release assay. Briefly target cells were incubated with Cr51 radionucleotide (Perkin Elmer, NEZ030S001MC) for one hour, washed and incubated with NK cells at increasing effector to target ratios, or in the absence of effectors for spontaneous release controls, in triplicate at 37C for 4 hours. After 4 hours, total experimental release controls were lysed with 1% octylphenoxypolyethoxyethanol. Plates were spun and the supernatant was transferred to a LumaPlate (PerkinElmer), dried overnight, and read in a gamma scintillation counter. Total lysis and spontaneous lysis controls were included. Percent specific lysis was calculated using the following formula: [(experimental release-spontaneous release)/(total release-spontaneous release)]*100.

### Western blotting

Cells were lysed in RIPA buffer (Thermo Cat# 89900) with 1X Halt Protease and Phosphatase Inhibitor cocktail (Thermo Cat# 78443). Cell lysates were separated by gel electrophoresis using a 4%-12% gradient cell (Thermo Cat# NW04120BOX) or 12% gel (Thermo Cat# NW00127BOX) and then transferred to a 0.2 *μ*m pore-size nitrocellulose membrane (Thermo Fisher Cat# LC2000). The membranes were blocked using 10% nonfat dry milk solution (Fisher Scientific Cat# 50-488-786) in PBS-Tween-20 0.1%, then incubated with primary antibodies in a 5% BSA solution. Membranes were imaged on the LiCOR CxL and analyzed using the ImageStudioLite software (version 5.2.5).

### Statistics

Statistical analyses were conducted using Prism 10.0.0 (GraphPad). All data show mean ± SEM. P ≤ 0.05 was considered significant. One-sample 2-tailed Student’s t test was used to compare the mean of an experimental condition to the control, normalized to one. One-way ANOVA was used to compare multiple conditions. Values were statistically non-significant when P>0.05. P values on graphs are represented as: * P *≤* 0.05, ** P *≤* 0.01, *** P *≤* 0.001, **** P *≤* 0.0001.

## Data sharing statement

Original datasets and protocols will be available for analysis by contacting. Proteomics data is available through the ProteomeXchange Consortium public database (https://www.proteomexchange.org/).

## ACKNOWLEDGEMENTS

We would like to thank the staff of the Columbia Stem Cell Initiative Flow Cytometry Core Facility, under the leadership of Michael Kissner, at Columbia University Irving Medical Center for their contributions to the work presented in this manuscript. These studies also used the resources of the Herbert Irving Comprehensive Cancer Center Flow Cytometry Shared Resources funded in part through Center Grant P30CA013696. This work was supported by grants to 1R01GM132604-01A1 to DE and R01AI137275 to EMM. The schematic illustration was created using biorender.com.

