## Supplemental Figures for "Replication stress in activated human NK cells induces sensitivity to apoptosis"

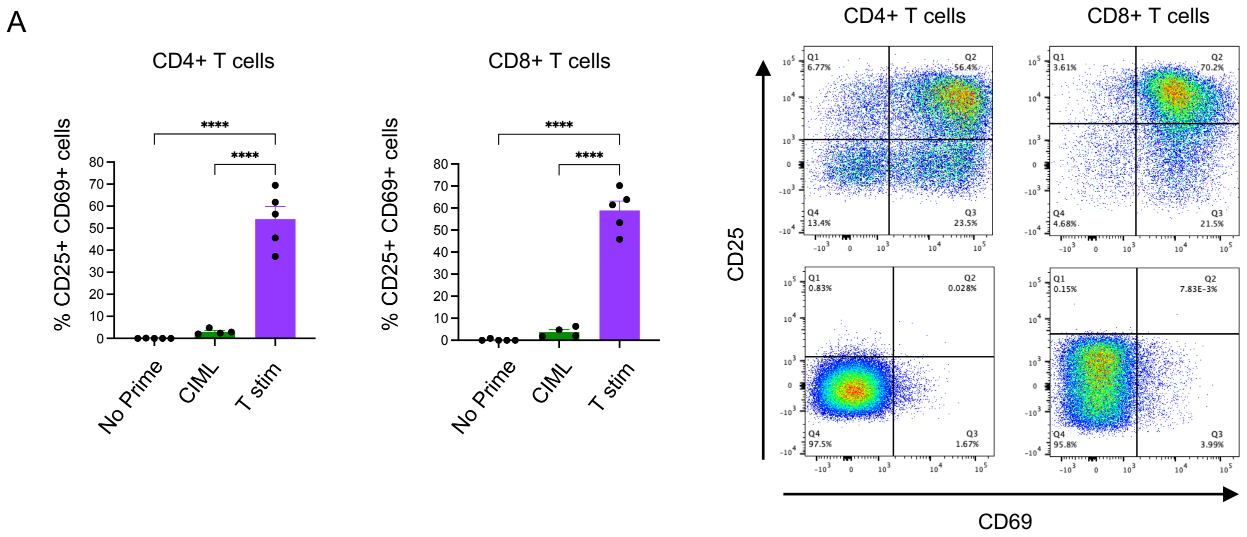


**Supplemental Figure 1: T cell activation.** (A) Percentage of activated CD25$^+$CD69$^+$ T cells. Gating is based on no stimulation controls for CD4$^+$ or CD8$^+$ cells, respectively.

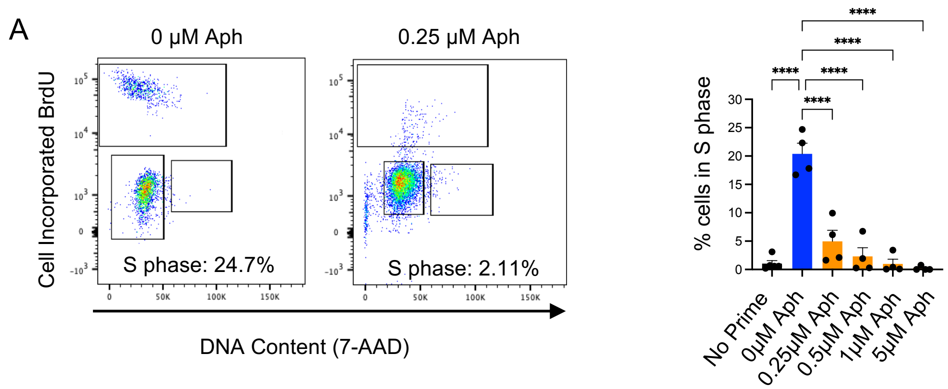


**Supplemental Figure 2: Cell cycle analysis during NK cell activation with replication stress.**

(A) Representative flow plots of cell cycle gating for CIML stimulation with and without low dose aphidicolin. Quantitative analysis of percentage of cells found in S phase. n=4 healthy donors One-way ANOVA calculated statistical significance.

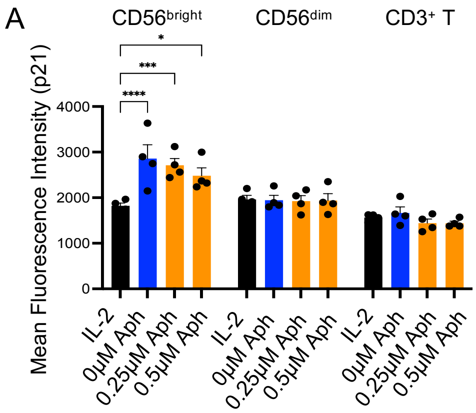


**Supplemental Figure 3: p21 analysis of activated NK vs T cells with and without replication stress.** (A) Fluorescence intensity of p21 was collected for CD56$^{bright}$ and CD56$^{dim}$ NK cells and CD3$^{+}$ T cells. n=4 biological replicates.

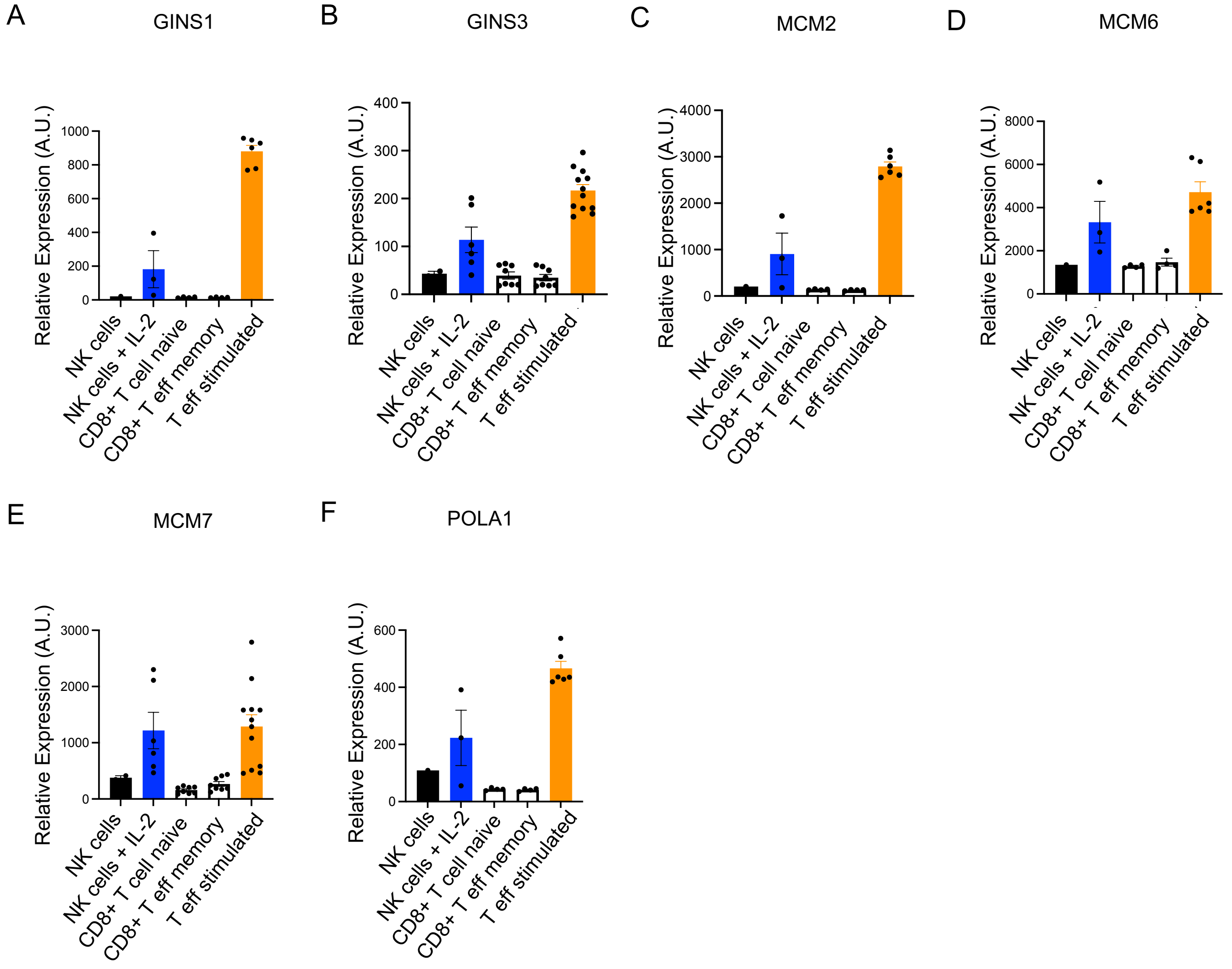


**Supplemental Figure 4: Gene expression of helicase and replication-related genes.** (A – F) Relative expression in arbitrary units for RNA abundance of given helicase proteins and replication genes was collected and analyzed from BioGPS. Significance was determined using a One-way ANOVA.
